# Exploration of the Link Between COVID-19 and Alcoholic Hepatitis from the Perspective of Bioinformatics and Systems Biology

**DOI:** 10.1101/2023.01.14.524034

**Authors:** Tengda Huang, Bingxuan Yu, Xinyi Zhou, Hongyuan Pan, Ao Du, Jincheng Bai, Xiaoquan Li, Nan Jiang, Jinyi He, Kefei Yuan, Zhen Wang

## Abstract

**Objective:** Severe acute respiratory syndrome coronavirus 2 (SARS-CoV-2) has been suggested to purpose threats to health of mankind. Alcoholic hepatitis (AH) is a life-threatening acute and chronic liver failure that takes place in sufferers who drink excessively. During the epidemic, AH has an increasing incidence of severe illness and mortality. However, for these two diseases, the intrinsic relationship of molecular pathogenesis, as well as common therapeutic strategies are still poorly understood.

**Methods:** The transcriptome of the COVID-19 and AH has been compared to obtain the altered genes and hub genes were screened out through protein-protein interaction (PPI) network analysis. Via gene ontology (GO), pathway enrichment and transcription regulator analysis, a deeper appreciation of the interplay mechanism between hub genes were established.

**Results:** With 181 common differentially expressed genes (DEGs) of AH and COVID-19 were obtained, 10 hub genes were captured. Follow-up studies located that these 10 genes typically mediated the diseases occurrence by regulating the activities of the immune system. Other results suggest that the common pathways of the two ailments are enriched in regulating the function of immune cells and the release of immune molecules.

**Conclusion:** This study reveals the common pathogenesis of COVID-19 and AH and assist to discover necessary therapeutic targets to combat the ongoing pandemic induced via SARS-CoV-2 infection and acquire promising remedy strategies for the two diseases.

## Introduction

SARS-CoV-2 can motive an acute respiratory sickness in humans, recognized as COVID-19(1). Up to 11 December 2022, extra than 645 million confirmed instances and over 6.6 million deaths have been reported by WHO(2). COVID-19 is a universal health problem, not only due to the fact of the rapid unfold of the virus from individual to individual, but also because of the massive and far-reaching impact on social life, the financial system and infrastructure(3). Recent findings indicate that disease severity is strongly associated with impaired immune response and comorbidity (4). At the equal time, enhancing the expression of ACE2, which is one of the viral host receptors, can block the entry of the virus and protect against infection through the novel coronavirus and potentially different coronaviruses(5). Severe COVID-19 frequently presents pathologically with pulmonary and extra-pulmonary organ dysfunction(6). Previous research suggested that the lungs are the organs most affected by the way of the novel coronavirus accompanied with a massive quantity of immune cell activation and the release of inflammatory factor(7, 8). Extra-pulmonary organs, on the other hand, have varying levels of tissue injury and inflammatory response, demonstrated as multi-organ dysfunction and systematic inflammatory reaction(9). Liver is the representative of extrapulmonary organs. Hepatocytes and cholangiocytes were observed to express the ACE2 receptor required for the virus to enter the cell(10). Others detected SARS-CoV-2 viral particles and positive stranded RNA with replication intermediates in hepatocytes with hepatocyte infection(11), indicating that SARS-CoV-2 also duplicates within hepatocytes. Since sufferers with COVID-19 and previously existing AH have elevated chances of severe disorders and death(12, 13), studies are necessary to perceive the interactions between AH and COVID-19.

Alcoholic liver disease (ALD) is the worldwide-existing disease which is the leading cause of alcohol-related deaths. Frequently, the first clinical presentation of ALD is AH, which is a lifethreatening acute and chronic liver failure that occurs in patients who continue to drink heavily (14). It is generally believed that alcohol has harmful effects on the intestine, which ultimately leads to liver inflammation through a variety of mechanisms. In particular, alcohol exerts direct toxicity to intestinal epithelial cells and reduces the expression of tight junction proteins, which enhance the permeability of the intestinal mucosa and allow pathogen-associated molecular patterns (PAMPs) such as lipopolysaccharides (LPS) to shift to the liver(15, 16). For AH, the effective treatment has not existed, and the only effective steroid drugs are not proven to have a long-lasting treatment effect(17).

Aiming to discover the co-pathogenesis of COVID-19 and AH, this study focused on the genes altered while diseases happen. Firstly, the datasets of SARS-CoV-2 and AH in the Gene Expression Omnibus (GEO) database have been analyzed and their DEGs were accessed, which supported further comparison to grab common DEGs. With the shared DEGs, the enrichment pathways and functions of these genes have been examined to recognize the process in which they participated. Besides, the relationship between all DEGs was shown with the aid of mapping PPI networks and gene-transcription factors (TF) interactions, and the hub genes with the highest level of interplay relationship was once screened from the shared DEGs. Follow on, biological role of these key genes was analyzed to explore their achievable mechanisms in disease progression. The sequential workflow we studied is shown in Fig. 1.

**Fig. 1.**
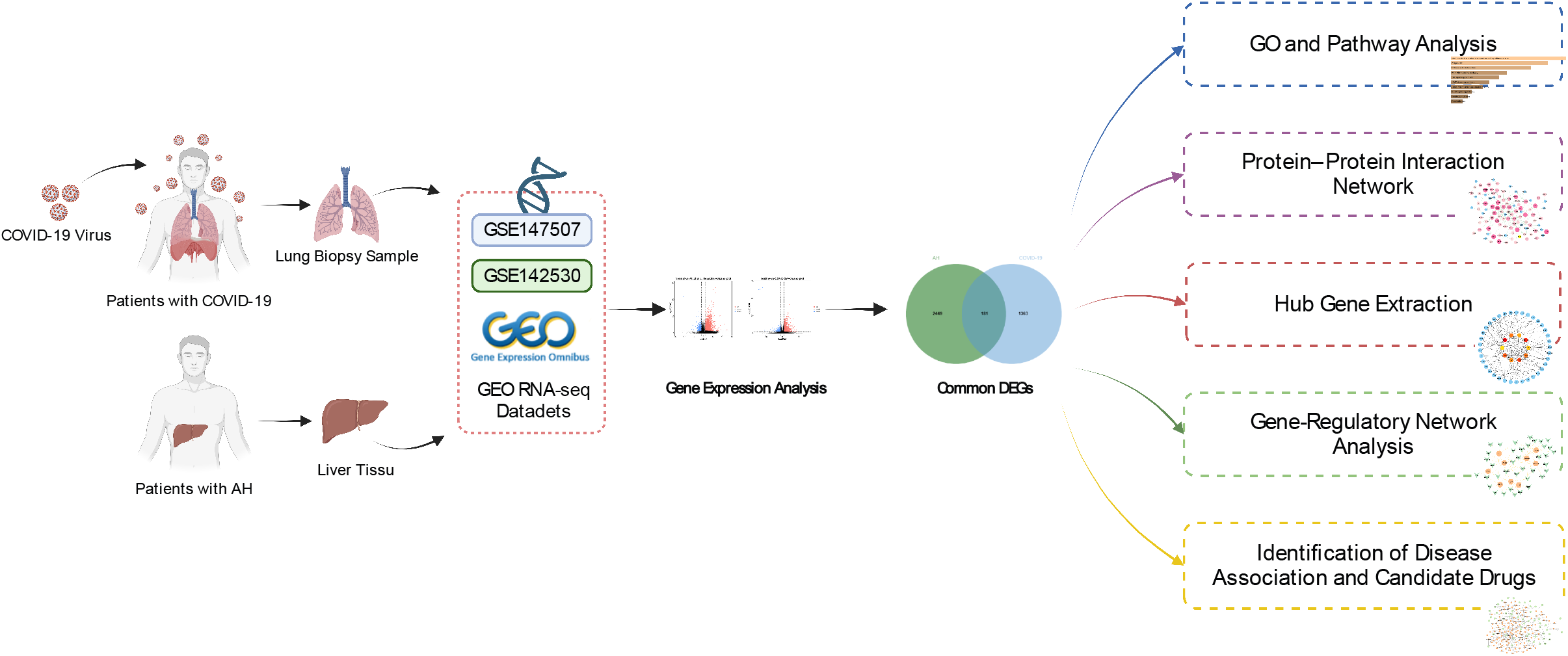
Schematic diagram of the overall general workflow for this study.

## Materials and Methods

### Data Source

All RNA-seq datasets for AH and COVID-19 in our research were collected from GEO (https://www.ncbi.nlm.nih.gov/geo/) database of National Center for Biotechnology Information (NCBI) (18). The dataset of SARS-CoV-2, GSE147507, is a transcriptome maps of lung biopsy in response to respiratory infection through the high-throughput sequencing Illumina NextSeq500 (Homo sapiens) platform, including 44 infection groups and 10 healthy controls. The data package for AH is GSE142530, which was processed by the Illumina HiSeq2000 (Homo sapiens) high-throughput sequencing system and included 10 alcoholic hepatitis samples and 12 healthy samples as controls.

### Identification of DEGs and Selection of Shared Genes

Obtaining the DEGs of AH and COVID-19 is critical to discovering the biological associations between them. Taking false discovery rate (FDR) < 0.05 and |log2 Fold Change | ≥1 as the screening conditions, R (version 4.0.1, https://www.R-project.org/) package DEseq2 (version 1.28.1)(19) with Benjamini-Hochberg correction was utilized to obtain DEGs of AH and 2019-nCoV respectively. Then intersect their DEGs to select shared genes by Jvenn (http://jvenn.toulouse.inra.fr/app/example.html)(20).

### GO and Pathway Enrichment Analysis

With the common DEGs of AH and SARS-CoV-2 detected, GO analysis considering biological process (BP), cellular component (CC) and molecular function (MF) were performed and their potential functions and pathways were figured out using Enrichr (https://maayanlab.cloud/Enrichr/), which is an all-around gene set enrichment web tool(21). After that, KEGG (Kyoto Encyclopedia of Genes and Genomes), WikiPathways, Reactome and BioCarta were used to classify the interactive networks of SARS-CoV-2 infections.

### PPI Network Analysis

Proteins expressed by mutual DEGs need to critically evaluate and integrate the interactions between them through PPI network analysis, which found direct and indirect associations in these genes(22). In this study, STRING (www.string-db.org)(version 11.5) for this analysis(23) was utilized to manage these shared DEGs. PPI network of common DEGs was constructed under the use of a combined score greater than 0.4. For a more intuitive visual effects and other experimental tests of the PPI network, Cytoscape (version.3.9.1) were chosen to plot our PPI network(24).

### Hub Gene Extraction

The top 10 hub genes in the PPI network were selected using Cytoscape plugin CytoHubba’s MCC method (a tool for sequencing and extracting central or possible target factors in biological networks primarily based on a number of network characteristics(25)) collected, at the same time, classifying the shortest accessible pathways of hub genes in accordance with the nearest rating traits of Cytohubba.

### Recognition of Hub DEGs Associated TFs and miRNAs

On the foundation of the hub genes, TF that regulate their transcription were found in the JASPAR (http://jaspar.genereg.net) of NetworkAnalyst platform, which is an open access resource for TF flexible models (TFFM) and TFs of massive species in 6 different taxonomic groups (26). At the same time, topological analysis with Tarbase and mirTarbase was performed to find out the miRNAs interacting with the target genes(27, 28). Via Cytoscape the interconnections between TF-genes and miRNA-genes were visualized. Screen out important ones, and the biological functions and characteristics were monitored.

### Gene-disease Association Analysis

Gene-disease associations were detected via NetworkAnalyst and DisGenet (http://www.disgenet.org/, a platform that integrates and standardizes statistics on disease-associated genes and varies from diversified sources(29)), which also identified other diseases associated with hub genes. The current version of DisGenet carries more than 24,000 ailments and traits, 17,000 genes and 117,000 genomic variations that can better help us in finding gene-related diseases(30). In this study, merely when the disease is related with at least two genes can it be shown on the gene-disease network.

## Result

### Recognition of Shared Transcriptional Signatures between COVID-19 and AH

In order to find the biological associations between AH and COVID-19, we found DEGs for two diseases and intersected them. Two volcano maps demonstrate DEGs for AH (Fig. 2A.) and COVID-19 (Fig. 2B.) diseases respectively. The results showed that 2,630 DEGs in AH (2,132 up-regulated DEGs and 498 down-regulated DEGs) and 1,544 DEGs were identified in COVID-19 (1,202 up-regulated DEGs and 342 down-regulated DEGs). 181 common DEGs between two diseases were extracted by Venn diagram (Fig. 2C), which has been shown to be the Transcriptional Signatures for AH and COVID-19.

**Fig. 2.**
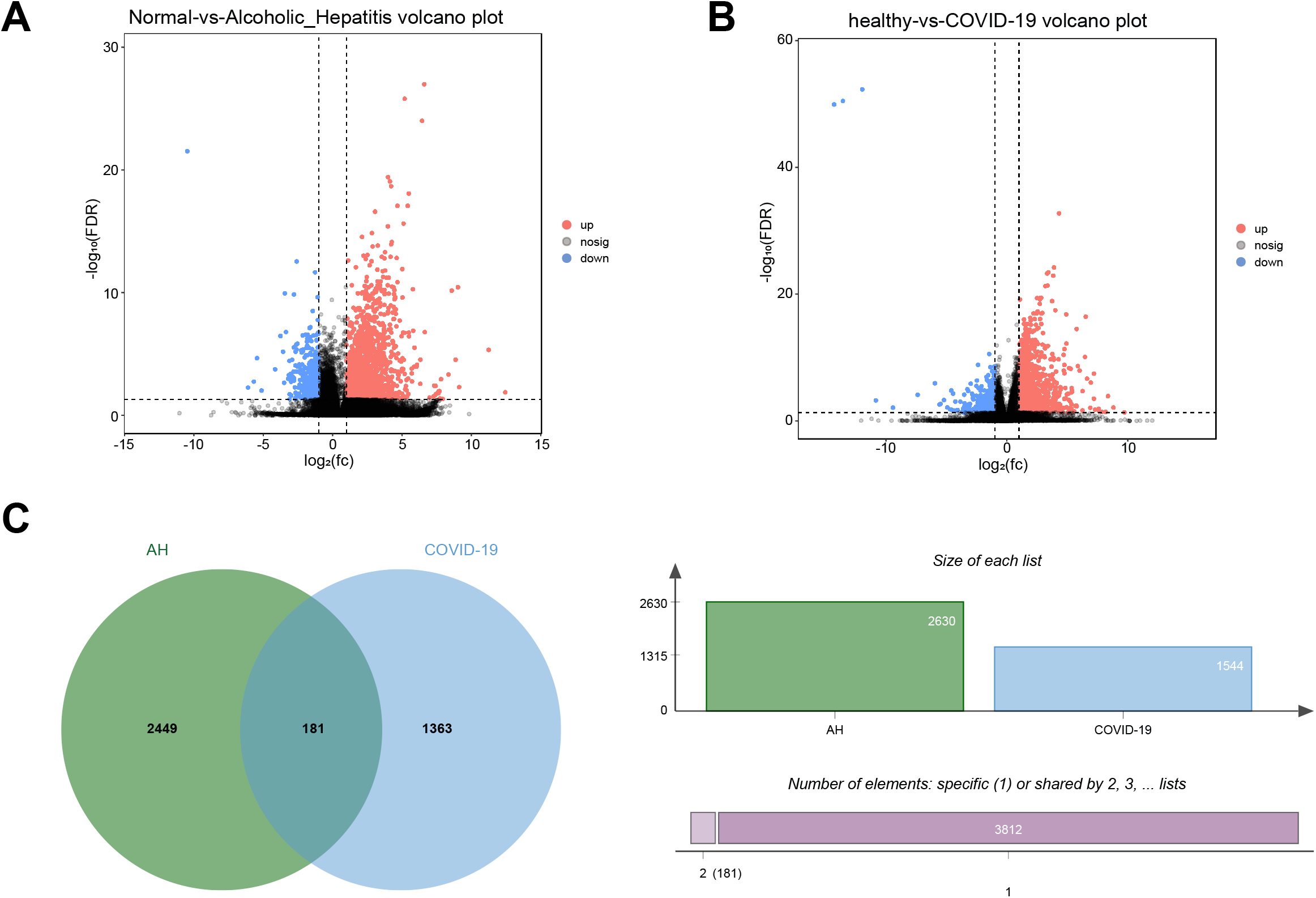
Volcano plots exhibit DEGs of (A) AH and (B) COVID-19. Red dots indicated up-regulated genes, blue dots indicated down-regulated genes and gray dots indicated non-DEGs. Figure C demonstrates that AH (green circle) and COVID-19 (bule circle) have 181 common DEGs in between.

### Functional-enrichment analysis identifies significant GO terms and pathways

In order to obtain the GO terms and the signaling pathways of common DEGs, we used the Enrichr tool to perform gene ontology analysis and pathway enrichment analysis. The results were displayed in Fig. 3 and Fig. 4.

**Fig. 3.**
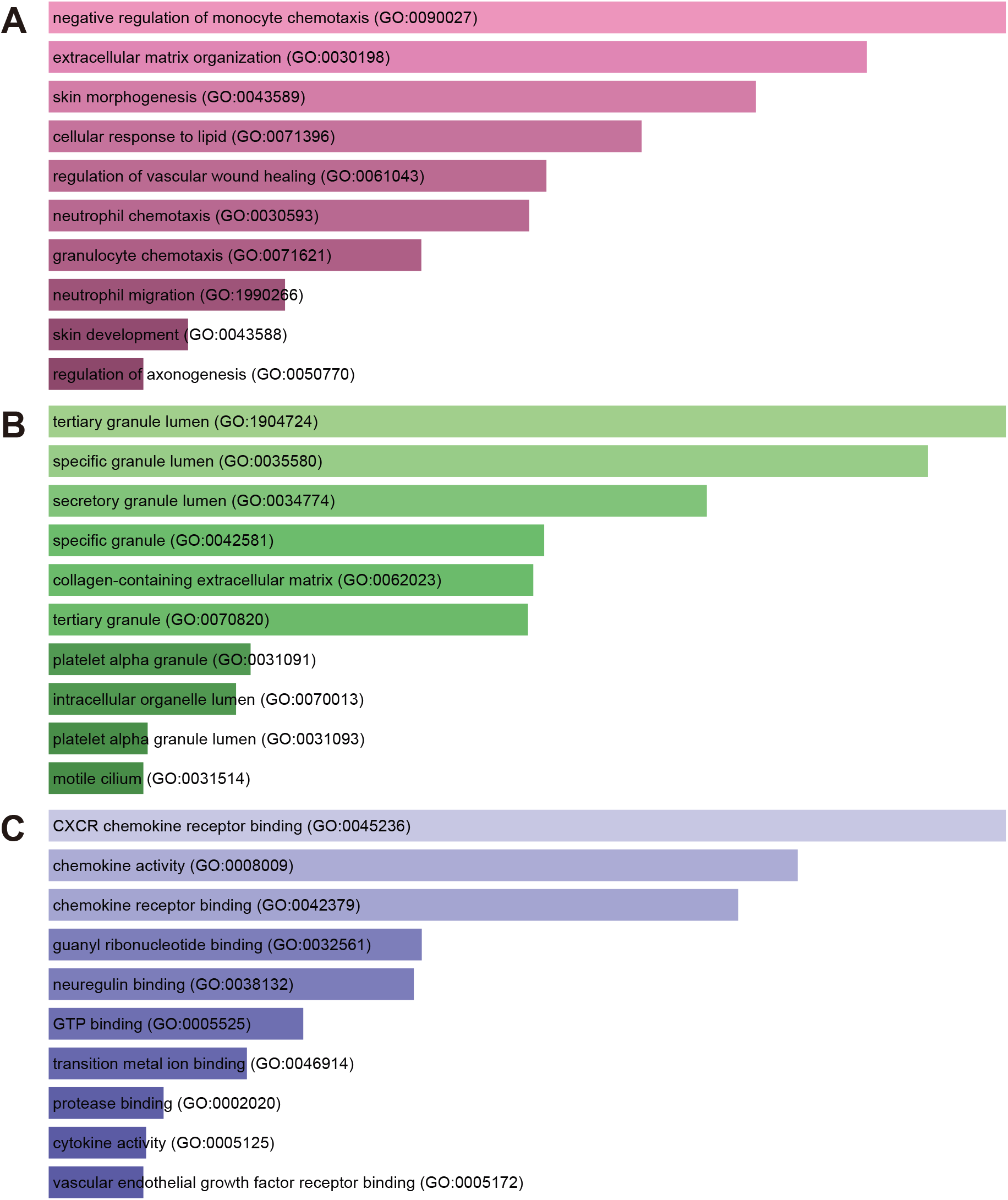
The bar graphs of the ontological analysis of the common DEGs between COVID-19 and AH. (A) biological processes; (B) cellular components; and (C) molecular functions.

**Fig. 4.**
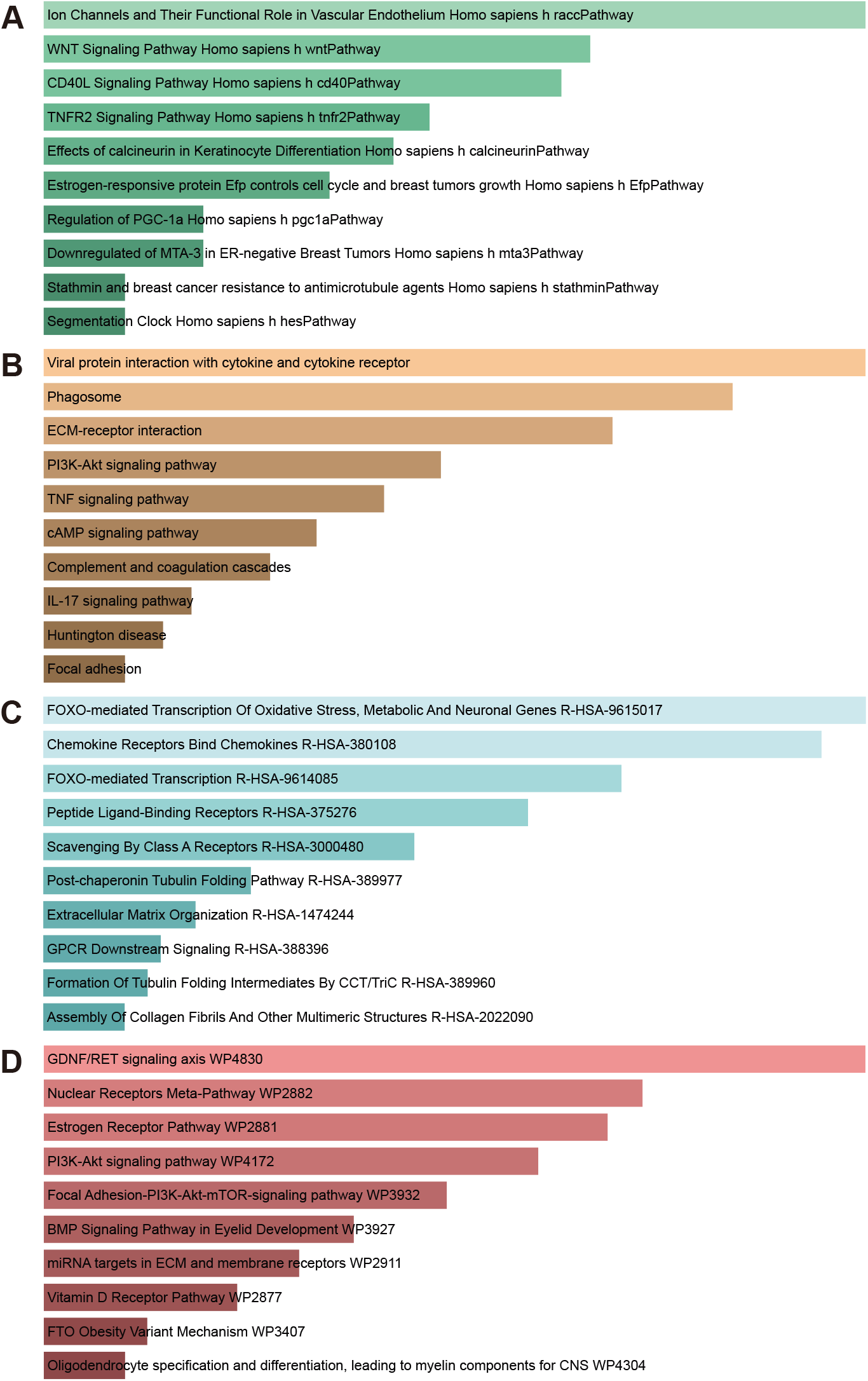
The bar graphs of pathway enrichment analysis of the common DEGs between COVID-19 and AH performed by (A) BioCarta 2016, (B) KEGG 2021 Human, (C) Reactome 2022, (D) WikiPathway 2021 Human.

BP (Fig. 3A.), CC (Fig. 3B.) and MF (Fig. 3C.) constitute the GO analysis. The top 10 GO terms of MF, BP, and CC were displayed in Table 1. BP found that common DEGs directed the function of macrophages and granulocytes in chemotaxis effects. Meanwhile, in the CC analysis, the formation or release of intracellular granular cavities were the main effects that shared DEGs generated. Similarly, MF found that pathway function is enriched in immune processes, including the regulation of immune molecule activity and mediating the binding of active molecules to receptors.

**Table 1.**
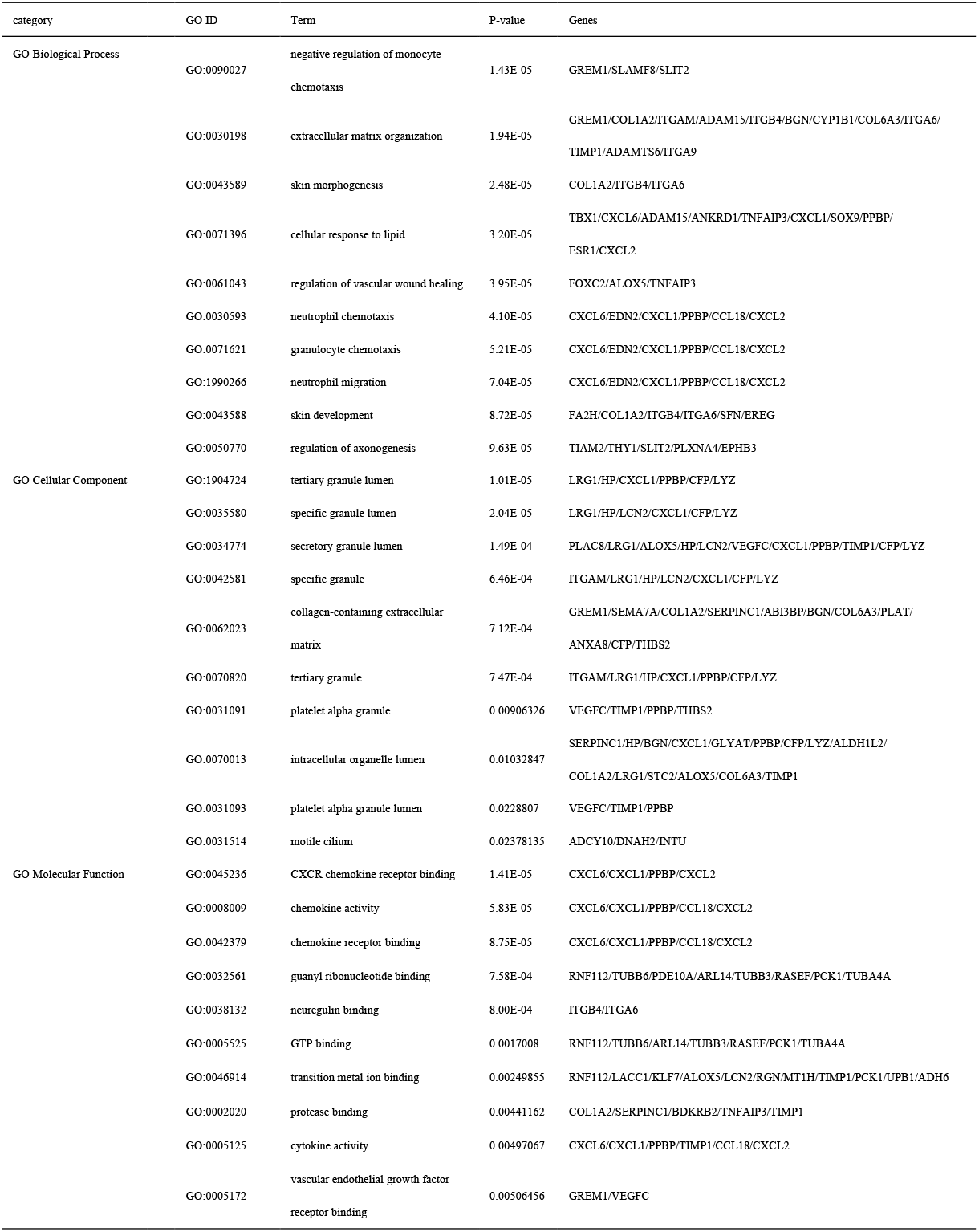
GO analysis of common DEGs among COVID-19 and AH.

In the results of pathway enrichment analysis, Fig. 4A, 4B, 4C and 4D represent the enrichment analysis results in four databases. Specific analysis revealed that BioCarta 2016 terms, such as WNT signaling pathway, CD40L signaling pathway and TNFR2 signaling pathway, have a strong connection with immune function. In addition, the results of KEGG 2021 Human also showed that common DEGs are connected through immunity process of ECM-receptor interaction, PI3K-Akt signaling pathway, TNF signaling pathway and IL-17 signaling pathway. As for Reactome 2022, similar results of immune process of FOXO-mediated Transcription and Chemokine Receptors Bind Chemokines were obtained. Furthermore, WikiPathway 2021 Human analysis suggested that pathway terms were mainly concerned with immune system, either. Table 2 shows the top 10 pathways for each of the four databases.

**Table 2.**
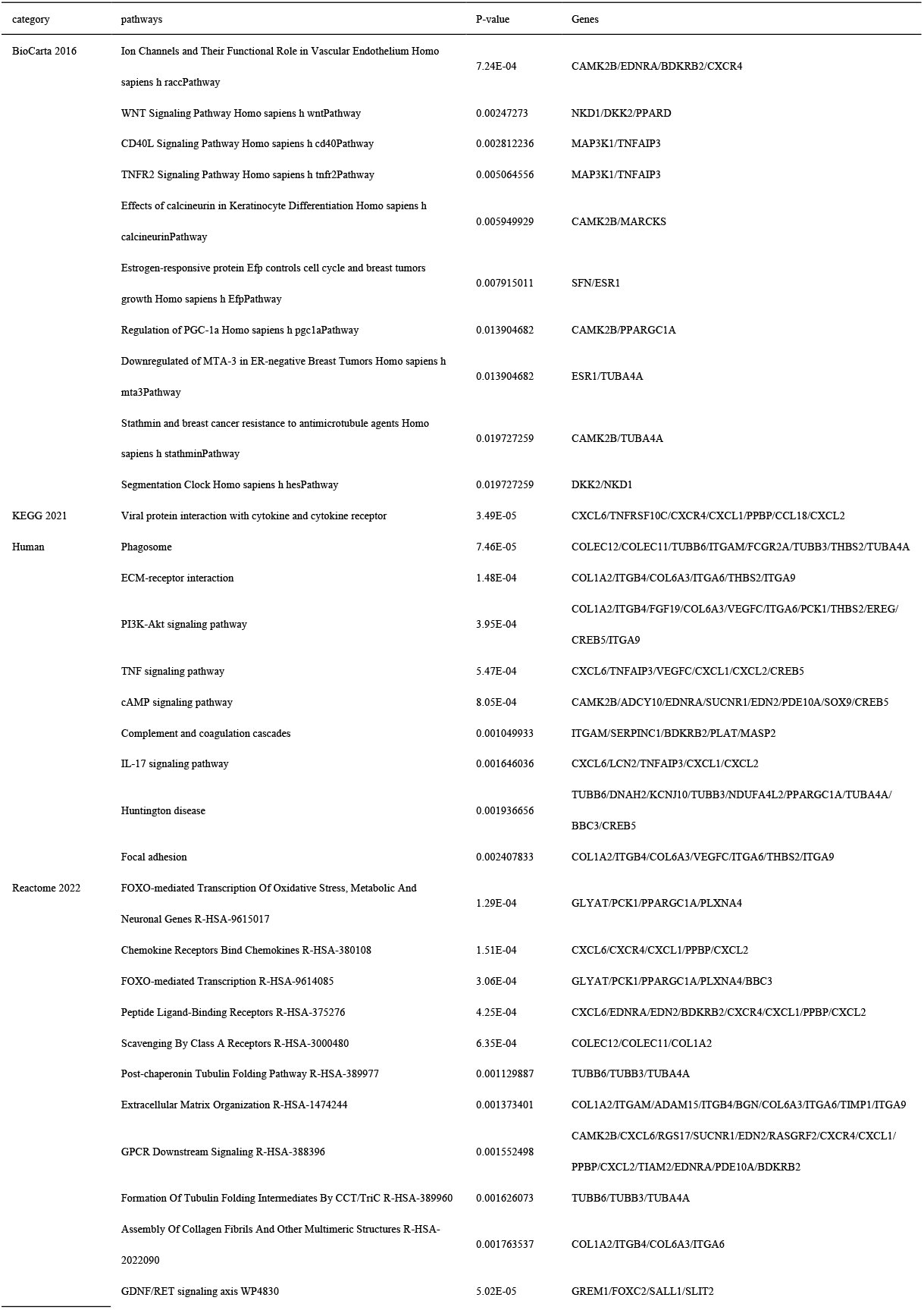
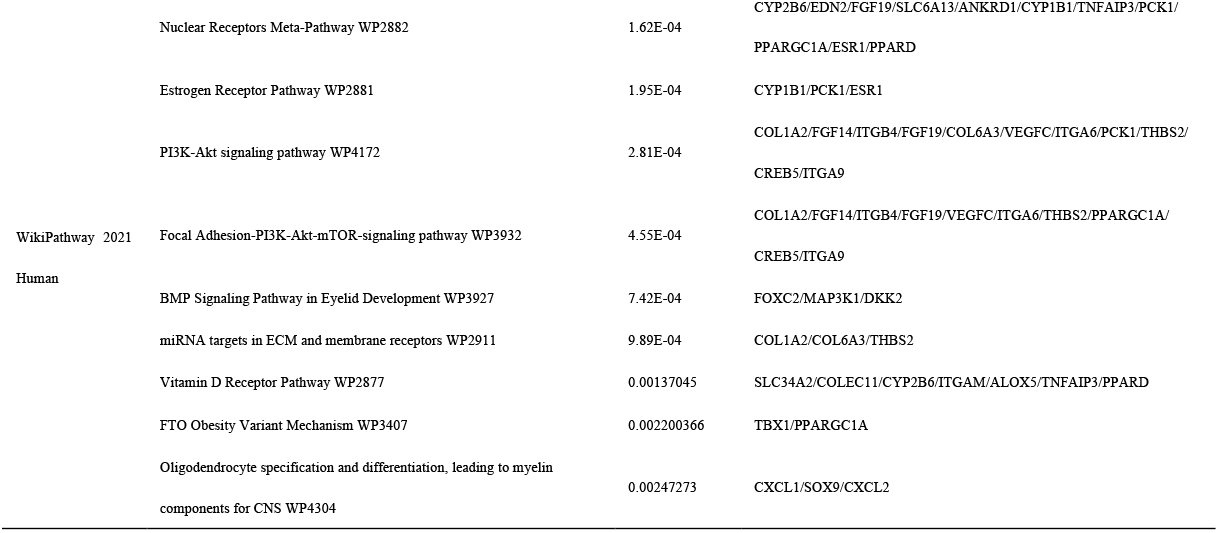
Pathway enrichment analysis of common DEGs among COVID-19 and AH.

### PPI Network Analysis and Recognition of Hub Genes

The relationship between interacting DEGs also requires PPI network analysis to evaluate and integrate. Once we have the hub protein, hub gene with high confidence could be obtained. Fig. 5 depicts the PPI network of shared DEGs between COVID-19 and AH, which consists of 103 nodes and 229 edges (nodes were considered as hub proteins). Meanwhile, the PPI topology table with nodality, intermediate centrality, stress centrality and close centrality is shown in Table 3. The top 10 common DEGs ranked by score, including CXCR4, COL1A2, ITGA6, ITGAM, TIMP1, THBS2, THY1, SOX9, COL6A3, CXCL1, were selected by the CytoHubba plug-in in Cytoscape after PPI network analysis and were considered to be the most influential hub genes. Specific information of hub genes is shown in Table 4. Since the hub gene is potential, a network of submodules, which consists of 47 nodes and 151 edges (nodes were considered as hub genes), was constructed with the help of the CytoHubba plugin to have a deeper understanding of reciprocal relationship between TFs and DEGs (Fig. 6).

**Fig. 5.**
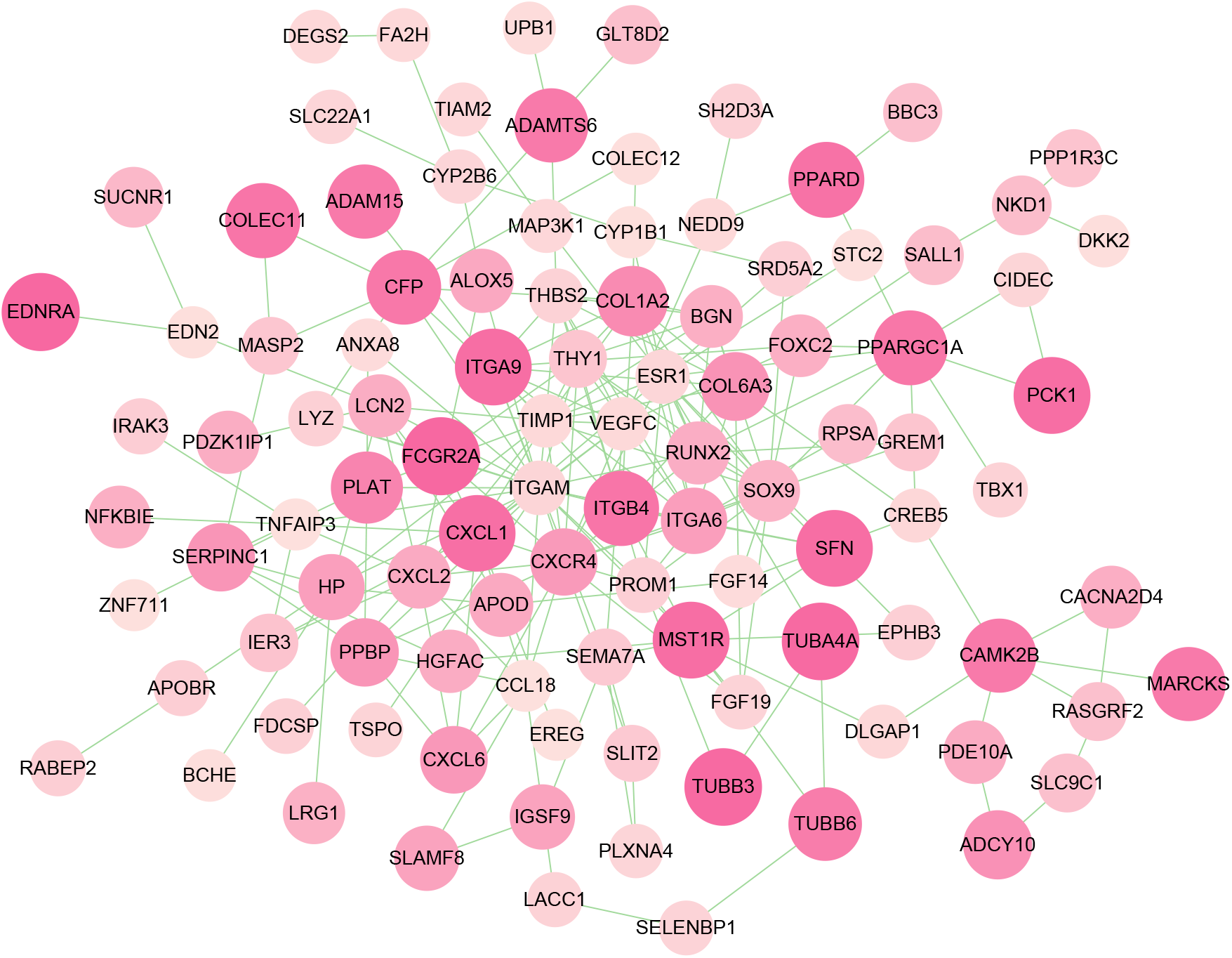
The PPI network of common DEGs among COVID-19 and AH. In the figure, the octagonal nodes represent DEGs and edges represent the interactions between nodes.

**Fig. 6.**
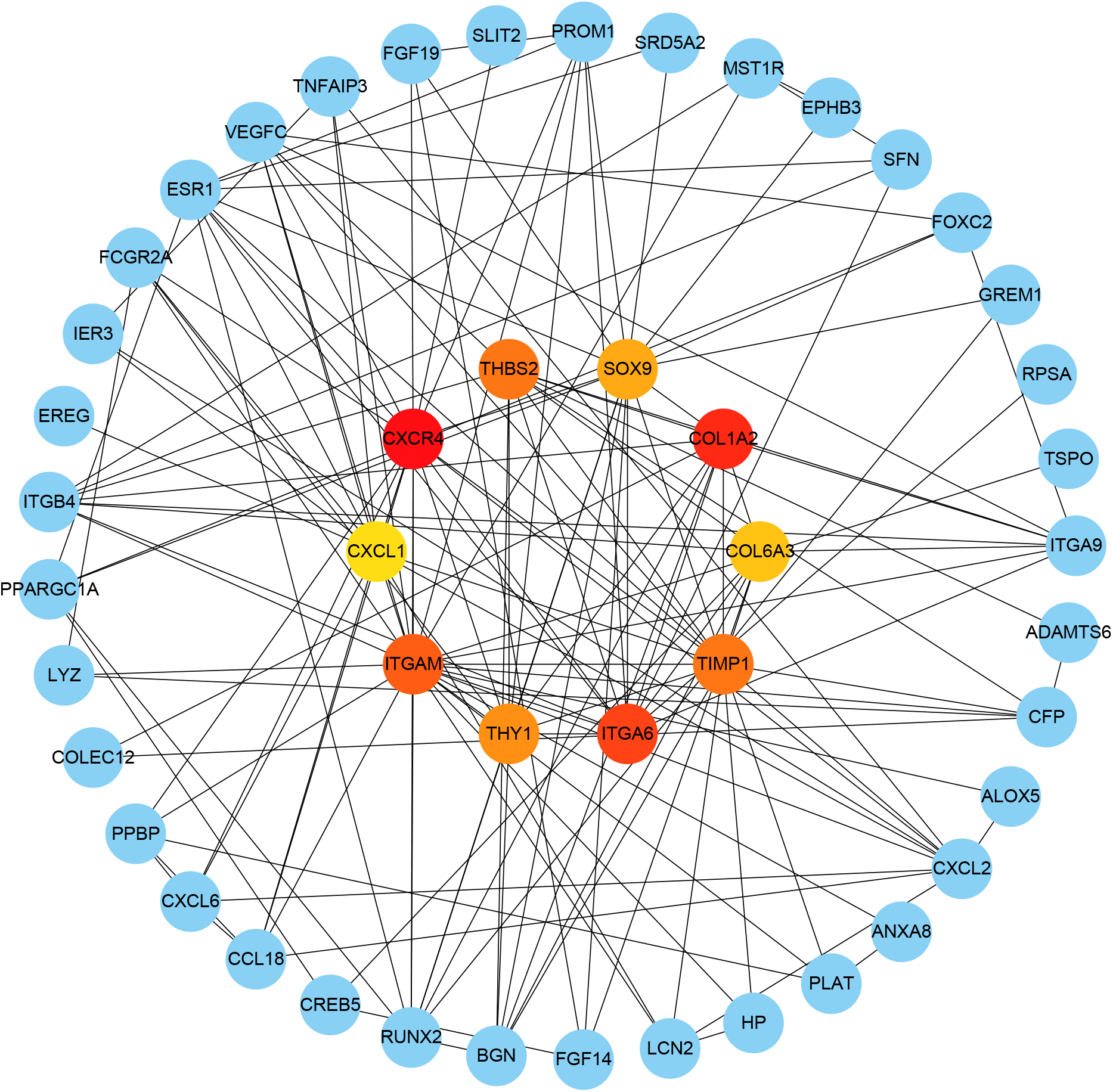
The hub gene was identified from the PPI network using the Cytohubba plug in Cytosacpe. Here, the colored central nodes represent the highlighted top 10 hub genes and their interactions with other molecules.

**Table 3.** Top 10 hub genes in network medium.tsv ranked by MCC method.

| Rank | Name | Score |
| --- | --- | --- |
| 1 | CXCR4 | 241 |
| 2 | COL1A2 | 229 |
| 3 | ITGA6 | 215 |
| 4 | ITGAM | 187 |
| 5 | TIMP1 | 182 |
| 5 | THBS2 | 182 |
| 7 | THY1 | 162 |
| 8 | SOX9 | 147 |
| 9 | COL6A3 | 146 |
| 10 | CXCL1 | 145 |

### Determination of regulatory signatures

In order to understand the commonality of the interaction between these Hub genes, we used the proteins and miRNA that regulate their transcription and translation as entry points through network analysis. Tarbase and Mirtarbase were used to identify the DEG-TFs interaction network (Fig. 8.). The orange circle represents the shared DEGs, while the green arrows represent the TFs. We consider Nodes with higher degrees as the pivots of the network. CXCR4, CXCL1, COL6A3, COL1A2 and ITGA6 have higher degree of expression in the TFs-gene interaction network, while TFs such as FOXC1, YY1, GATA2, SREBF1 and FOXL1 are higher than others (Fig. 8.). Similarly, the correlation between hub genes and miRNA is shown in Fig. 9. The blue box represents miRNA, while the red oval represents hub genes. The figure clearly shows that THBS2, SOX9, THY1, COL1A2 and TIMP1 are the hub genes of the network, which are most closely related to miRNAs. In addition, prominent hub miRNAs were detected in the miRNAs-gene interaction network, namely hsa-mir-1-3p, hsa-mir-26b-5p, hsa-mir-29b-3p, hsa-mir-105-5P and hsa-mir-4781-5P. The interaction of these common DEGs with TFs and miRNAs may assist hub genes in regulating the occurrence of disease.

**Fig. 7.**
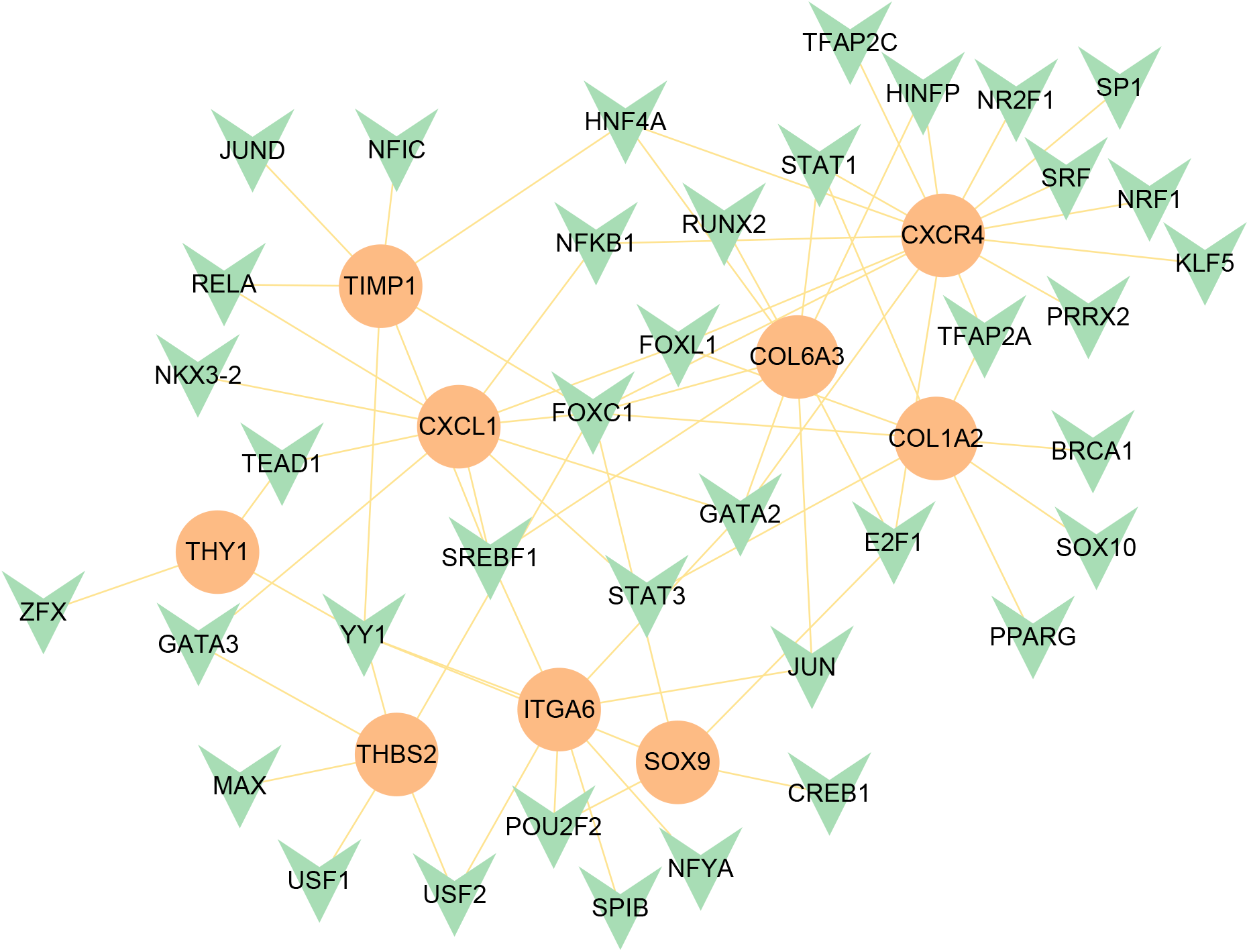
The Network Analyst created an interconnected regulatory interaction network of DEG-TFs. The orange circle represents the shared DEGs, while the green arrows represent the TFs.

**Fig. 8.**
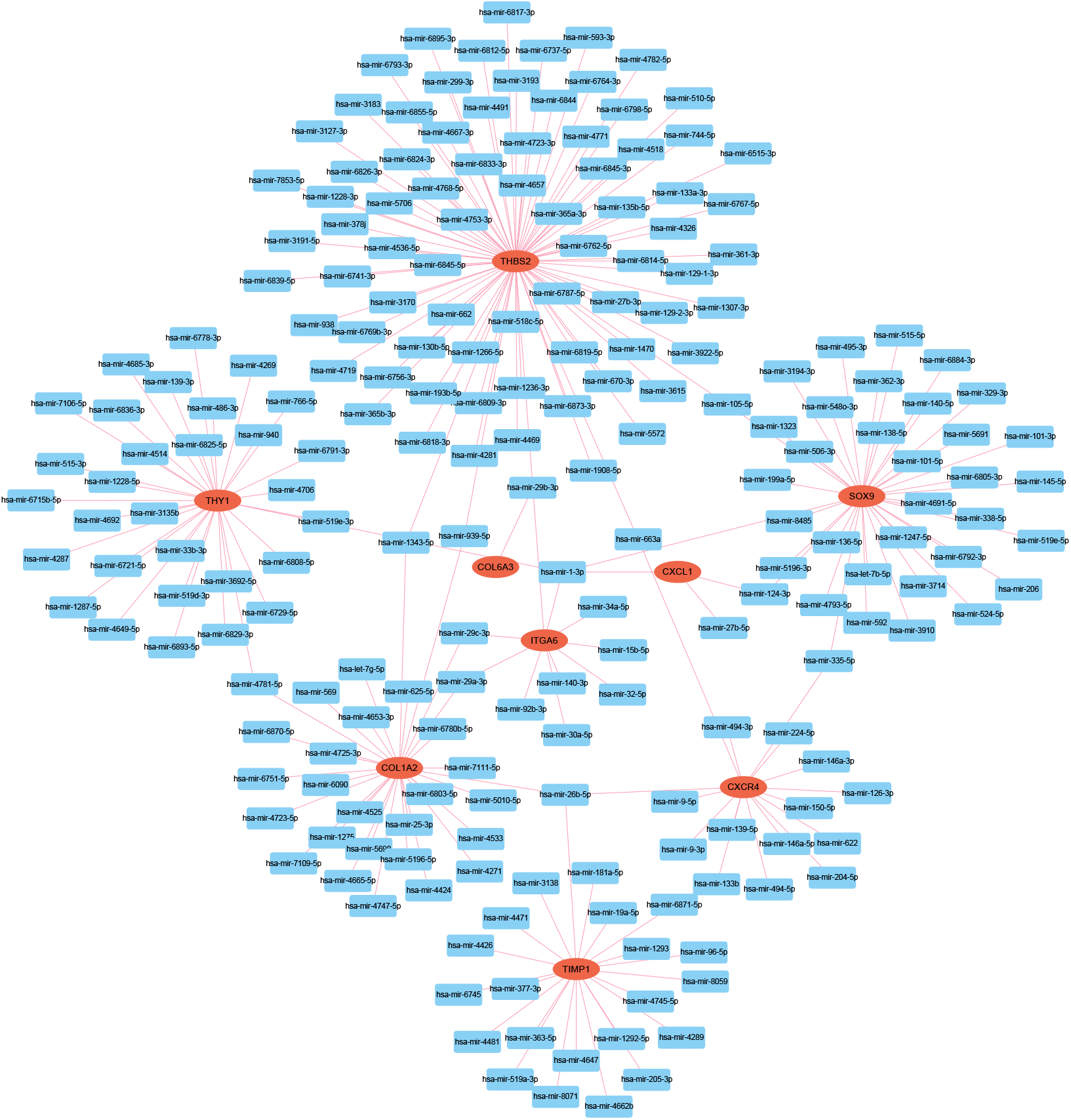
The interconnected regulatory interaction network of DEGs-miRNAs. The blue box represents miRNA, while the red oval represents hub genes.

**Fig. 9.**
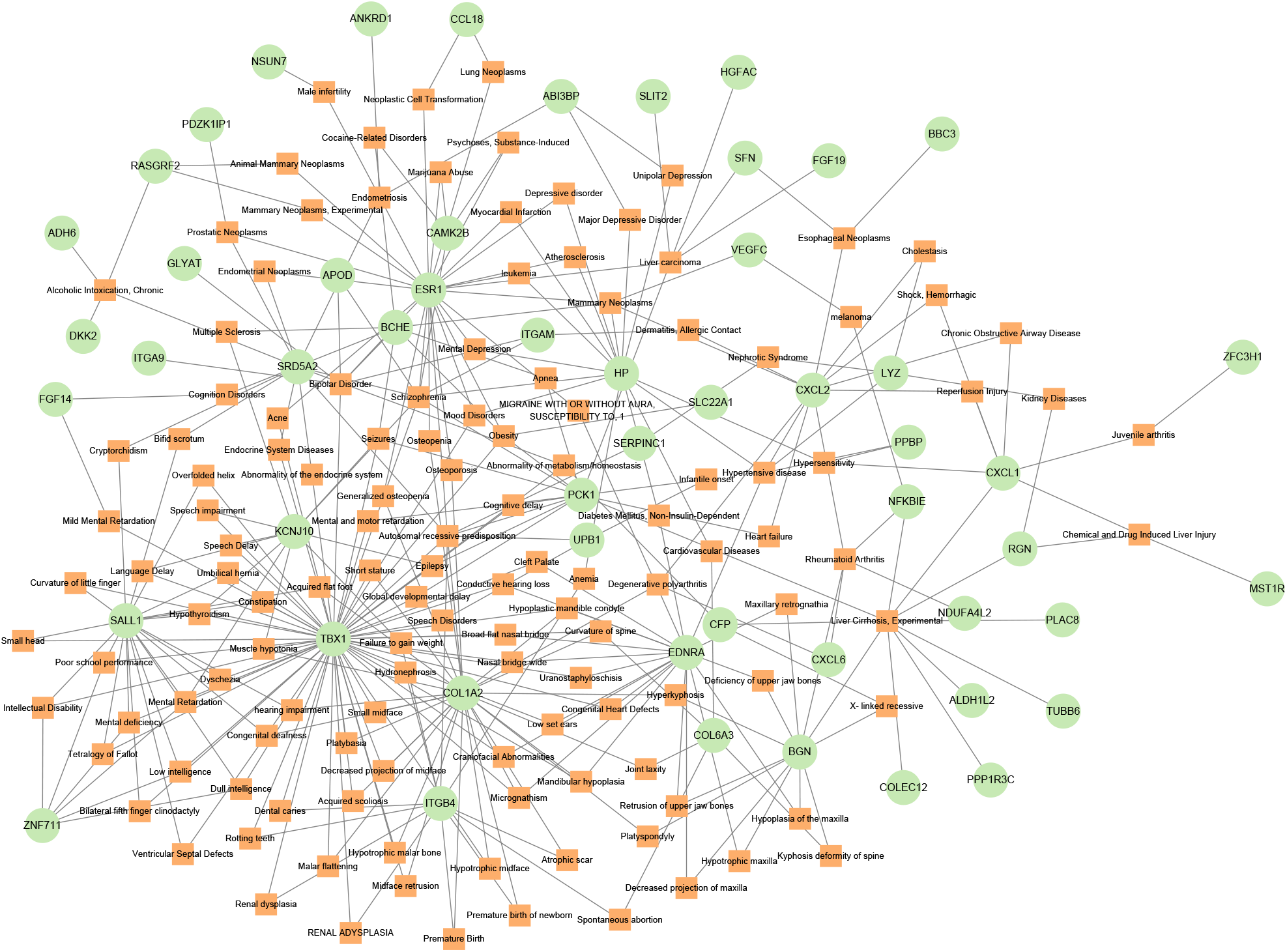
The gene-disease association network represents diseases associated with mutual DEGs. The disorder depicted by the square node and also its subsequent gene symbols is defined by the circle node.

### Identification of Disease Association

Therapeutic design strategies for diseases are beginning to reveal the relationship gene-disease relationships(31). From NetworkAnalyst, it was noticed that the abnormality of metabolism/homeostasis/endocrine were most relevant to our reported central genes. The associations between genetic diseases were shown in Fig. 9.

## Discussion

With the gradual deepening of the research on the COVID-19, it has become a consensus that there may be some association between it and AH. A study reported that the rate of acute chronic liver failure precipitated with acute severe alcoholic hepatitis significantly raised during the COVID-19 pandemic (12). Another study reported that sufferers with chronic liver disease including AH, are susceptible to severe COVID-19(13). Through epidemiological studies between AH and COVID-19, it has also been found that patients with severe COVID-19 have an accelerated incidence of abnormal liver function while patients with liver disease are identified to be at high risk for severe COVID-19 infection, and AH may also be a hazard factor for severe COVID-19 and adverse outcomes(32). On this basis, COVID-19 and AH have been speculated to have quite rich links at the genetic level.

GO is a gene regulation context based on the generic theoretical model that accelerates genes and their internal connections(33). Using Enrich R of the GO database as the basis for ontology analysis, three types of GO analysis were carried out. For BP, the 181 common DEGs were mainly enriched in the chemotaxis regulation of monocytes and granulocytes, which is part of the complex immune response to SARS-CoV-2. In the MF analysis, it is still emphasized that these common genes have more commonality in the expression of chemokines and their receptors. Chemokines are important inflammatory mediators in the antiviral immune response and are responsible for infiltrating immune cells into infected lungs(34). Studies on lung type II lung cells (A549) have shown that the open reading framework (ORF) 7 in the genome of the new coronavirus induces the production of the chemokine CCL2 that promotes the chemotaxis of monocytes and reduces the expression of the chemokine IL-8, which recruits neutrophils. ORF7 increases monocytes infiltration, reduces the number of neutrophils, and may have a specific effect on immunological changes in tissues after infection(35). Similarly, for the results of a study of AH suggest that in primary biliary cirrhosis (PBC), there is a functional monocyte disturbance which seems to be primary and may arise at the cellular level and is the way that SARS-CoV-2 could anabasis(36). The results of CC analysis also fully support the above research results, which are reflected in the formation and release of granulocytes granule cavities. Neutrophils, as “professional” phagocytes, possess a large number of antibacterial “weapons” in their granules, allowing them to destroy pathogens during phagocytosis. During the degranulation process, bactericidal enzymes can also be released from the cell(37).The mechanism of this process leading to SARS-CoV-2 exists in the process of neutrophil extracellular traps (NETs) modified by bactericidal proteins in granules and cytoplasm, which indicates that we can disrupt the vicious cycle of hepatocellular inflammation in patients with new coronary pneumonia by activating neutrophils and promoting the formation of NETs(6).

Pathway analysis is the best way to reflect the organism’s response through internal changes(33). BioCarta human pathway includes WNT, CD40L and TNFR2 signaling pathways, and regulation of PGC-1a. Here, the researchers found that WNT’s response to β-catenin-mediated inflammatory activity in alveolar macrophages contributed to the host’s acute incidence of COVID-19. Mechanistically, WNT treatment promotes β-catenin-HIF-1α interactions and glycolysis-dependent inflammation, while inhibiting mitochondrial metabolism, thereby inhibiting AM proliferation, i.e., enhanced expression of HIF-1α is caused by the WNT signaling pathway and causes macrophage inflammation in COVID-19 patients(38). During the occurrence of AH, the researchers found that WNT/β-catenin signaling was negatively correlated with the expression of Foxo3A through rat model studies, and reduced steatosis, cell damage and apoptosis in ALD rats. Activation of WNT/β-catenin signaling inhibits Foxo3A-induced apoptosis by upregulating serum/glucocorticoid-regulated kinase 1 (SGK1). In addition, pharmacological recovery of WNT/β-catenin signaling reduces the progression of ALD in vivo(39). The KEGG human analysis also mentions viral protein interaction with cytokine and cytokine receptor, PI3K-Akt, TNF signaling pathway and IL-17 substance. Among them, the IL-17 pathway has been observed to upregulate expression in the serum of COVID-19 patients in multiple studies and has been speculated to induce a cytokine storm to promote inflammation, while its inhibitors have been observed to attenuate or even prevent cytokine storms(40). This process has also been reported in the pathogenesis of AH, through intermediate monocytes expansion, functional activation, induction of CD4 T cell IL-17 expression, and enrichment in the liver of AH patients to obtain greater phagocytic capacity(41). Reactom pathway and Wikipathway analysis also found that the common mechanism of AH and COVID-19 exists in pathways such as mediating chemotaxis effects, complementing the first two analyses.

Using 181 DEGs, a PPI network was constructed based on an in-depth understanding of protein biology. According to the PPI network and hub gene extraction, CXCR4 has more reciprocal actions with other genes and is probably the most important gene between COVID-19 and AH. The findings echo those of others who have found that deadly COVID-19 also manifests as an escalation of activation of CXCR4^+^ T cells in bystander lungs and enhances SARS-CoV-2-specific T effector responses, while reducing CXCR4-mediated homing may help recovery from severe disease(42). Similarly, the researchers found in mice with AH that Mallory-Denk body (MDB) formation correlated with MyD88-dependent TLR4/NFκB pathway regulation in AH formed by the NFκB-CXCR4/7 pathway(43). The common pathways of other hub genes are also important. For example, ITGAM, a defensive factor expressed at some stage in inflammatory damage, encodes integrin-CD11b which is expressed mostly at the surface of macrophages and is concerned with adherence, migration, and cell-mediated cytotoxicity. CD11b mediating thrombosis in new coronary pneumonia proves that ITGAM plays a vital role in thrombosis in patients infected with SARS-CoV-2(44, 45). ITGAM is also identified as a marker of liver damage and is differentially expressed by macrophages during the disease process(46). In addition, TIMP1 can also be a promising non-invasive prognostic biomarker for COVID-19 patients, promising for selecting cures focused on matrix metalloproteinase pathways in patients(47). It additionally contributes to fibrogenesis in hepatitis C virus (HCV) infection and in alcohol-induced liver disease (ALD), which could lead to AH(48). Therefore, recognized hub genes can be viewed potential biomarkers and become the novel drug target after biological insight in COVID-19 is confirmed.

For purpose of recognizing how shared DEGs alter COVID-19 (or AH) at the transcriptiome level, the interactions amongst TFs, miRNAs and hub genes have been identified by means of internet tools. The results showed that TFs (FOXC1, YY1, GATA2, SREBF1 and FOXL1) were associated with CXCR4, CXCL1, COL6A3, COL1A2 and ITGA6 and miRNAs (hsa-mir-1-3p, hsa-mir-26b-5p, hsa-mir-29b-3p, hsa-mir-105-5P and hsa-mir-4781-5P) were closely regulated with THBS2, SOX9, THY1, COL1A2 and TIMP. In previous bioinformatics analyses, Ahmed, Islam, and Lu Lu et al. found FOXC1, YY1, GATA2, and FOXL1 are significant TFs for COVID-19 (6, 49, 50). Pharmacological studies have also found that SREBF1, as a key regulator of lipid-metabolizing enzymes connected closely with the procession of AH, is the transcription factor most activated by several effective antiviral drugs in vitro and may become A pivot therapeutic spot for both AH and COVID-19 (51, 52). In addition, hsa-mir-1-3p, as a well-known antiviral miRNA, may induce activation or inhibit and affect gene expression of linkage genes, and cooperates with TFs to regulate autophagy hub genes (53, 54) and hsa-mir-26b-5p destroys mRNA target sites or mRNA functions through strong hybridization with ACE2, which is also considered to be a potential therapeutic target for COVID-19 (55). Although many preceding research have proven that these TFs and miRNAs may additionally equip vital treatment effects, these research statistics still require similar further experiments to affirm their effectiveness and reliability.

Gene-disease networks based on hub genes were utilized to gain a deeper understanding of the relationship between genes and disease for COVID-19 and AH. For example, the study found that the hub genes, which influence the occurrence of AH and COVID-19, also affect disorders of metabolism, homeostasis and endocrine in the body. Metabolic disorders, including those of iron, lipid and glucose (diabetes), have been reported in other network meta-analyses(56–58). Another research reported that up to 50% of the sufferers who have died from COVID-19 had metabolic and vascular problems and there are many direct hyperlinks between COVID-19 and the metabolic and endocrine systems(59).

## Conclusions

Transcriptome data for novel coronary pneumonia and AH and normal controls were downloaded from public databases, then were utilized to find DEGs and finally used to obtain 181 common DEGs for both diseases. To further understand how these common DEGs act on the disease process, gene ontology analysis and pathway enrichment analysis were performed by Enrichr tools. The results show that they are mainly involved in chemotaxis effects, cell granules and granular cavities and immune-related pathways and functions. Next, to clarify the interaction between these shared DEGs, it was need to perform PPI network analysis, obtain 10 hub genes and map gene-TFs interaction with gene-miRNA interaction at the level of transcription factors and miRNAs. Analysis of AH and COVID-19 suggests a way to identify infections from a variety of diseases, so it is feasible to mitigate the risk of COVID-19 in patients with AH. Because there were not many studies on the hazard elements and linked diseases of COVID-19, the results above assist people to fully understand the SARS-CoV-2. Currently, there are vaccines available to prevent coronavirus. Though these vaccines are not effective in some cases, especially against different variants of the novel coronavirus. Still, the scientific world is focused on developing a more effective vaccine to treat COVID. Therefore, transcriptional analysis was implemented to detect mutual pathways and molecular biomarkers that contribute to understanding the link between SARS-CoV-2 and AH. Overall, the genes identified could become new therapeutic targets for the development of new coronary pneumonia vaccines.

## Acknowledgements

We are most grateful for GEO databases providing their platform and thanking contributors who have uploaded their valuable datasets. Additionally, we appreciated Core Facility of West China Hospital for their technique support.

